# Genome-resolved metagenomics reveals seasonally defined life strategies in hypersaline sediments

**DOI:** 10.64898/2026.10.19.760023

**Authors:** Habib Ullah, Ramganesh Selvarjan, Maggie C. Y. Lau Vetter

## Abstract

**Background:** Hypersaline sediments are long-standing models for microbial adaptation and are increasingly regarded as reservoirs of novel biosynthetic potential, including surface-active compounds. Inferences about biosurfactant production in such communities have rested largely on taxonomic affiliation, targeted single-gene surveys or metabolite detection, rather than on genome-resolved evidence that the pathways are encoded. The seasonal restructuring of these communities has also rarely been resolved at genome level.

**Results:** We sequenced summer and winter sediments from the hyperarid Qaidam Basin on the northern Tibetan Plateau, recovering 44 metagenome-assembled genomes and 1,468 biosynthetic gene clusters from 2.98 Gbp of assembly. Evaporative and cryoconcentration processes raised winter porewater chloride by two to three orders of magnitude relative to summer. The community switched between osmoadaptation strategies. Summer sediments were dominated by salt-in haloarchaea with acidic proteomes and rhodopsin phototrophy, whereas winter sediments were dominated by salt-out bacteria with near-neutral proteomes and winter-enriched compatible-solute biosynthesis. The salt-in bacterium *Salinibacter* occupied the expected intermediate position, providing an internal control. The winter surface-active response was envelope-associated rather than secreted, comprising phospholipase-derived lysophospholipids, desaturase-mediated membrane remodelling, substitution of phospholipids by phosphorus-free ornithine and sulfoquinovosyl lipids, and capsular exopolysaccharide export. Dedicated secreted biosurfactant pathways were not detected. Of 69 NRPS clusters, only one carried the starter condensation domain diagnostic of lipopeptide assembly, and it matched no characterised reference. Profile-based, read-level and orthology-based searches independently recovered no complete lipopeptide, rhamnolipid or trehalolipid pathway. Apparent biosurfactant genes in 40 of 44 genomes resolved to housekeeping homologues upon domain verification, and an independent catalogue of 1,773 regional genomes reproduced the same absence.

**Conclusions:** Surface activity in this hypersaline community is retained at the cell envelope rather than secreted, and dedicated biosurfactant biosynthesis is not a community-level trait. Because both taxonomic affiliation and gene-name annotation overestimate this capacity, genome-resolved verification is required before biosynthetic potential inferred from extreme environments is treated as real.

## Introduction

Hypersaline environments rank among the most physiochemically extreme habitats on Earth, and the microbial communities that inhabit them have long served as models for understanding the limits of life and the molecular basis of adaptation (Gunde-Cimerman et al. 2018; Oren 2008). Hypersaline sediments in particular combine high and fluctuating salinity with steep redox and nutrient gradients, supporting taxonomically and functionally distinctive communities dominated by halophilic archaea and specialized bacteria (Gunde-Cimerman et al. 2018; Oren 2008; Andrei et al. 2012). Beyond their value as adaptation models, such communities are increasingly recognized as reservoirs of novel biosynthetic potential: the selective pressures of extreme habitats are thought to favor the production of specialized metabolites, including surface-active and membrane-stabilizing compounds, that may have both ecological and biotechnological relevance (Waschulin et al. 2022). Yet the genome-resolved functional ecology of hypersaline sediments, which organisms carry which adaptive capacities, and how these are distributed across the community, remains comparatively underexplored, particularly in hyperarid basins such as the Qaidam Basin.

Coping with osmotic stress is the defining physiological challenge of hypersaline life, and microorganisms meet it through two broadly distinct strategies (Oren 2008; Roeßler and Müller 2001). In the salt-in strategy, employed by most halophilic archaea and a few specialized bacteria, molar concentrations of potassium chloride are accumulated in the cytoplasm, and the entire proteome is adapted to function at high internal ionic strength, an approach detectable as a genome-wide excess of acidic surface residues (Oren 2008). In the salt-out strategy, characteristic of most halophilic and halotolerant bacteria, the cytoplasm is kept low in salt and osmotic balance is achieved by synthesizing or importing organic compatible solutes such as ectoine, glycine betaine, and trehalose (Gunde-Cimerman et al. 2018). These strategies carry very different genomic and energetic approaches, and the balance between them within a community is a fundamental descriptor of how that community is organized. How this balance shifts over time, however, has rarely been resolved at the genome level.

This temporal dimension is critical because many hypersaline environments are strongly seasonal. Evaporative concentration, temperature, and water availability vary widely between seasons, imposing a moving target of combined osmotic, thermal, and desiccation stress (Gunde-Cimerman et al. 2018). Most metagenomic studies of such environments however represent single time points, leaving the seasonal restructuring of community composition and function largely uncharacterized. Whether communities track seasonal change by shifting between osmoadaptation strategies, and how their broader metabolic and biosynthetic capacity responds, is a question that genome-resolved, multi-season sampling is well positioned to address.

Among the adaptive traits proposed to emphasize life in extreme environments are surface-active, amphiphilic lipids, molecules that partition to interfaces and modulate the interaction between cells and their physicochemical surroundings (Bagheri et al. 2025; Citarasu et al. 2021). This broad class involves both secreted biosurfactants and bioemulsifiers and cell-associated amphiphiles such as membrane phospholipids, their remodeling products, and secreted exopolysaccharides. Secreted lipopeptide biosurfactants in particular have attracted attention for their roles in stress tolerance, surface colonization, and nutrient access, and for their biotechnological potential, and prior amplicon- and culture-based work has suggested that halophilic taxa may be enriched in such producers (Citarasu et al. 2021; Fariq and Yasmin 2020). However, inferences about biosurfactant production in extreme communities have largely rested on the taxonomic affiliation of community members, on targeted single-gene surveys, or on the detection of surface-active metabolites, rather than on direct, genome-resolved evidence that the biosynthetic pathways are present. This is an experimental gap that leaves open which amphiphilic-lipid strategies are actually encoded and by whom. Genome-resolved evidence carries its own difficulty: the committed enzymes of biosurfactant pathways belong to large families whose other members serve unrelated primary-metabolic roles, so annotation by gene name alone routinely assigns biosurfactant identity to housekeeping homologues.

Here we address these gaps in a seasonally sampled hypersaline sediment community from the Qaidam Basin, combining genome-resolved metagenomics of summer and winter sediments with our previous 16S rRNA amplicon profiling and metabolomics (Ullah et al., under review in *Environmental microbiome*). Motivated by the hypothesis that surface-active amphiphilic lipids contribute to microbial adaptation in this environment, we set out to test this hypothesis directly at the level of biosynthetic and metabolic genes, while placing it within a whole-community functional and taxonomic context. To ensure that functional and abundance inferences reflected the entire community rather than the small fraction of contigs recovered into genomes, we quantified abundance over the complete assembly at the genome, gene, and biosynthetic-cluster levels, and, because the archaeal share of the community changes markedly between seasons. This approach reveals a seasonal shift between salt-in and salt-out osmoadaptation strategies and identifies the community’s surface-active-lipid response.

## Materials and Methods

### Study site and sample collection

The Qaidam Basin (36°39′N, 96°29′E), on the northern Tibetan Plateau (approximately 2,700–3,200 m elevation) (Jiang et al. 2022), is a hyperarid endorheic basin widely recognised as a terrestrial analogue for ancient Martian lacustrine environments (Liu et al. 2025). It experiences annual precipitation below 50 mm, with parts of the western basin receiving as little as 15–20 mm, intense ultraviolet irradiation with 220–320 sunshine hours per month, (Kong et al. 2018), extensive evaporite mineralogy (halite and gypsum)(Z.-K. Li et al. 2025), seasonal temperature extremes ranging from a monthly average of −10.07 °C in winter to +16°C in summer (Kong et al. 2018), and pronounced salinity gradients driven by evaporative concentration (Anglés and Li 2017).

Sediment samples were collected from the Nuomuhong floodplain in the hyperarid western Qaidam Basin during two contrasting seasons: winter (January 2023) and summer (July 2024). Four sediment samples were selected for shotgun metagenomic sequencing based on microbial community patterns from a 16S rRNA amplicon study (Ullah et al., under review in *Environmental microbiome*): two summer samples (C3, C6) and two winter samples (H2, H6), chosen to capture seasonal dynamics of biosurfactant biosynthetic potential.

Samples were collected with sterile tools and gloves, transported under cold conditions, and stored at −80°C until processing. The full site description, sampling design, and physicochemical characterisation are reported in our previous study (Ullah et al., under review in *Environmental microbiome*). For context, sediment pH, moisture, electrical conductivity, soil N/P/K, and porewater ion concentrations (ion chromatography, Dionex ICS-6000, Thermo Scientific) for the four metagenome samples are shown in Table 1.

**Table 1.**
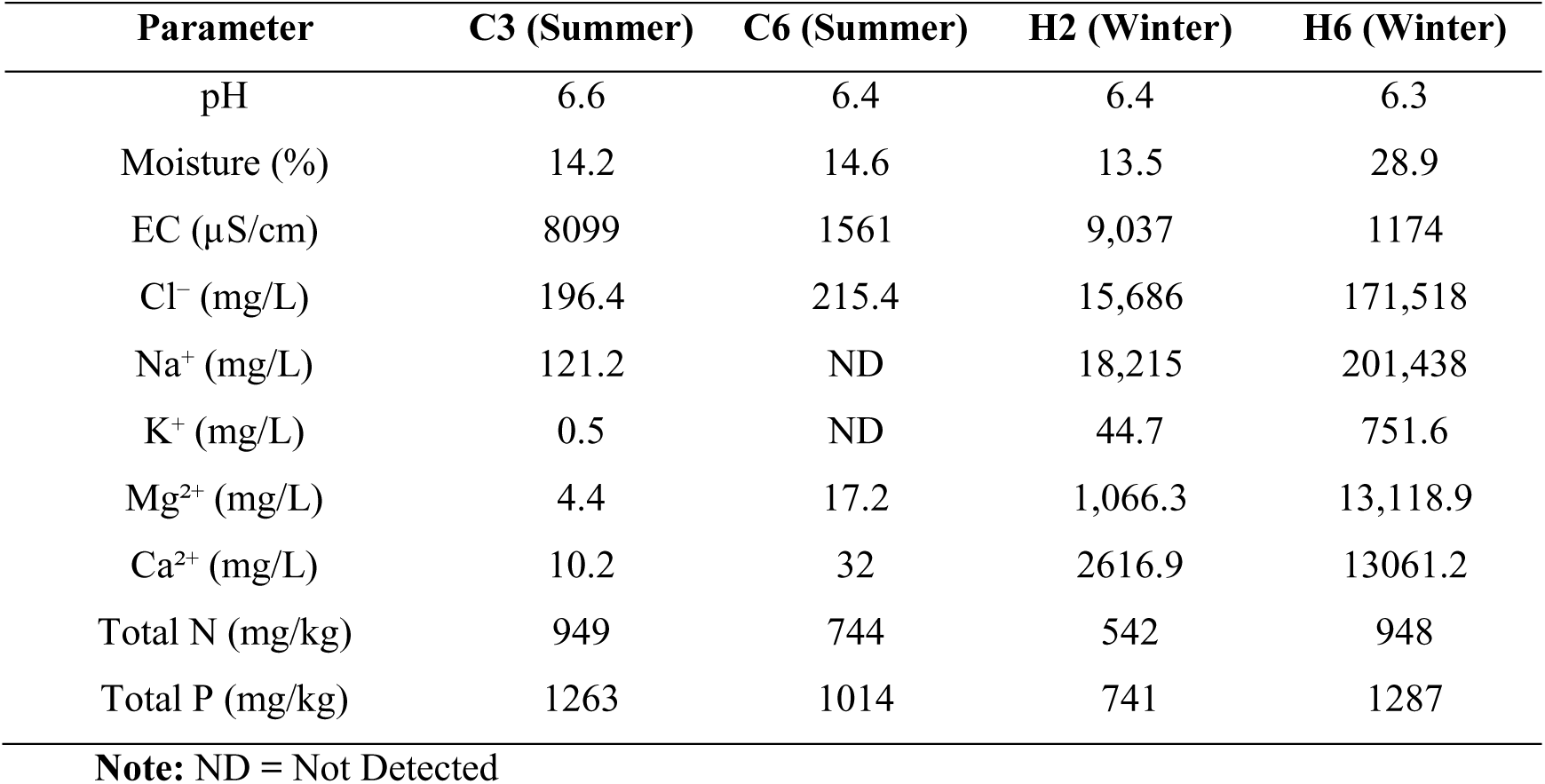
Physicochemical parameters of Qaidam Basin sediments across seasons.

| Parameter | C3 (Summer) | C6 (Summer) | H2 (Winter) | H6 (Winter) |
| --- | --- | --- | --- | --- |
| pH | 6.6 | 6.4 | 6.4 | 6.3 |
| Moisture (%) | 14.2 | 14.6 | 13.5 | 28.9 |
| EC (µS/cm) | 8099 | 1561 | 9,037 | 1174 |
| Cl<sup>-</sup> (mg/L) | 196.4 | 215.4 | 15,686 | 171,518 |
| Na<sup>+</sup> (mg/L) | 121.2 | ND | 18,215 | 201,438 |
| K<sup>+</sup> (mg/L) | 0.5 | ND | 44.7 | 751.6 |
| Mg<sup>2+</sup> (mg/L) | 4.4 | 17.2 | 1,066.3 | 13,118.9 |
| Ca<sup>2+</sup> (mg/L) | 10.2 | 32 | 2616.9 | 13061.2 |
| Total N (mg/kg) | 949 | 744 | 542 | 948 |
| Total P (mg/kg) | 1263 | 1014 | 741 | 1287 |

**Note:** ND = Not Detected

### DNA extraction and metagenomic sequencing

Total genomic DNA was extracted from 2 g of homogenised sediment per sample using an SDS-CTAB bead-beating protocol optimised for high-salinity sediments, as described in our previous study. Briefly, sediments were washed to remove soluble salts, followed by cell lysis in CTAB buffer using 300-µm glass beads and enzymatic treatments with lysozyme and proteinase K. DNA was purified by phenol:chloroform:isoamyl alcohol extraction and isopropanol precipitation.

Paired-end libraries were prepared and sequenced by Magigene Biotechnology Co., Ltd. (Shenzhen, China). Demultiplexed reads with adapter removed were provided by the sequencing facility (Table 2).

**Table 2.** Sequencing, quality-filtering, and assembly statistics per sample.

| Sample | Season | Raw reads | Raw Q30 (%) | Filtered reads | Filtered Q30 (%) | Retained (%) | Contigs ≥500 bp | Assembly (bp) | Max contig (bp) | Mapped reads | Mapping (%) | Number of BGCs |
| --- | --- | --- | --- | --- | --- | --- | --- | --- | --- | --- | --- | --- |
| C3 | Summer | 222,160,230 | 96.45 | 220,559,458 | 96.94 | 99.28 | 1,254,200 | 1,259,279,669 | 176,123 | 114,066,400 | 51.72 | 598 |
| C6 | Summer | 155,482,880 | 95.61 | 152,441,250 | 96.50 | 98.04 | 560,702 | 515,369,301 | 115,170 | 49,880,246 | 32.72 | 220 |
| H2 | Winter | 257,887,258 | 96.26 | 255,084,626 | 97.07 | 98.91 | 964,023 | 1,000,481,265 | 219,772 | 113,409,596 | 44.46 | 549 |
| H6 | Winter | 289,345,920 | 97.12 | 286,974,502 | 97.72 | 99.18 | 214,502 | 204,555,446 | 342,375 | 77,933,923 | 27.16 | 101 |
| Total | NA | 924,876,288 | 95.61–97.12 | 915,059,836 | 96.50–97.72 | 98.94 | 2,993,427 | 2,979,685,681 | NA | 355,290,165 | 27.16–51.72 | 1,468 |

**Note:** NA = Not Applicable

### Read processing and assembly

Reads were quality-filtered with fastp (Chen et al. 2018) (minimum Q20, automatic paired-end adapter detection). Putative human reads were removed by alignment to GRCh38 human reference genome (Ensembl release 102) using Bowtie2 v2.5.4 (Langmead and Salzberg 2012), retaining pairs in which both mates failed to map (SAMtools v1.9) (H. Li et al. 2009).

Each sample was assembled independently with MEGAHIT v1.2.9 (D. Li et al. 2015) under default metagenomic parameters (-t 56). Contigs shorter than 500 bp were removed with seqkit v2.13.0, and all downstream analyses were performed on this filtered contig set. For biosynthetic gene cluster detection, antiSMASH was additionally restricted to contigs ≥1,000 bp (--minlength 1000).

### Metagenomic binning, genome quality, and taxonomy

MAGs were recovered with MetaWRAP v1.3.2 (Uritskiy et al. 2018) using MetaBAT2, MaxBin2 and CONCOCT, consolidated through the MetaWRAP bin-refinement module (--min-completion 50 --max-contamination 10). Completeness and contamination were assessed with CheckM2 v1.1.0 (Chklovski et al. 2022). 44 MAGs meeting medium-to-high-quality thresholds were retained. Taxonomy was assigned with GTDB-Tk v2.3.0 (Chaumeil et al. 2022) against the GTDB release r214 reference database. To avoid cross-sample contig-name collisions during genome-resolved attribution, MAG-to-GTDB assignments were resolved in a sample-aware manner from the per-sample GTDB-Tk outputs.

### Open reading frame prediction and functional annotation

Protein-coding open reading frames (ORFs) were predicted from filtered contigs with Prodigal v2.6.3 in metagenomic mode (-p meta). ORFs were functionally annotated with eggNOG-mapper v2.1.12 against the eggNOG v5.0.2 database, using DIAMOND for homology searches. KEGG Orthology (KO), CAZy family, Pfam domain, and preferred gene-name fields were retained for downstream functional, abundance, and taxonomic analyses. Annotation was completed for all four metagenome assemblies.

### Biosynthetic gene cluster detection and novelty assessment

Biosynthetic gene clusters (BGCs) were identified with antiSMASH v7.1.0 (Buchfink et al. 2021) under relaxed detection stringency (--strictness relaxed) to recover both divergent and fragmented clusters. Full and cluster-level HMM scanning, TIGRFAM annotation, active-site prediction, and MIBiG v3.1 and ClusterBlast were enabled. Contigs below 1,000 bp were excluded from BGC detection (--minlength 1000). For lipopeptide assignment, antiSMASH KnownClusterBlast comparisons to MIBiG v3.1 were interpreted at the gene level: a cluster was accepted as a candidate lipopeptide only when the genes matching a characterized lipopeptide reference included its multi-modular synthetase. Matches resting solely on shared fatty-acid-supply genes (short-chain acyl-CoA synthetase, acetyl-CoA carboxylase) or on the two-gene KnownClusterBlast minimum were treated as non-specific and excluded.

To assess sensitivity to contig fragmentation, detection was repeated after filtering contigs to a 1,500-bp minimum (∼1.5× the mean assembly N50). BGCs were grouped into gene cluster families with BiG-SCAPE v1.1.9 (Navarro-Muñoz et al. 2020) in hybrid glocal mode (--mix - -mode glocal) at cosine cutoffs of 0.30, 0.50 and 0.70; cluster-family assignments are reported at a 0.50 cut-off. Families showing <0.50 cosine similarity to all MIBiG entries (Adnan, n.d.) were classified as lacking a characterized homologue. Because most clusters were truncated at contig boundaries (as indicated by antiSMASH contig-edge annotation), the proportion of novel families is reported as an upper bound.

### Quantification of taxonomic and functional abundance

Abundance was quantified over the complete assembly (all binned and unbinned contigs) so that all measures reference the total community rather than the recovered genomes alone. Metagenome-assembled genomes (MAGs) accounted for approximately 6% of assembled contigs, and genome-level abundances are therefore reported as a minority fraction of the community. For each sample, quality-filtered reads were mapped to the corresponding full assembly with Bowtie2 in end-to-end sensitive mode (--end-to-end --sensitive), and alignments were sorted and indexed with SAMtools. Between 27% and 52% of quality-filtered reads mapped to the corresponding assembly (C3 51.7%, C6 32.7%, H2 44.5%, H6 27.2%), of which 84–88% satisfied the identity and alignment-length thresholds applied for coverage estimation. Whole-community abundance measures therefore reference the assembled fraction of the community, which is substantially larger than the ∼6% of contigs recovered into genome bins. Abundance was estimated at three complementary levels.

Contig, MAG, and BGC coverage. Contig and BGC abundance were calculated as trimmed-mean coverage (10% trimming at each tail) with CoverM v0.7.0 (23) in contig mode (--min-read-percent-identity 95 --min-read-aligned-percent 75). Per-MAG abundance was obtained by aggregating the coverage across contigs belonging to each bin. Each antiSMASH BGC was assigned the coverage of its host contig, with the unbinned fraction retained explicitly.

### Gene-level functional abundance

From the same alignments, read pairs were counted per ORF using featureCounts (Subread v2.1.1) (-p --countReadPairs -M --primary) based on Prodigal derived gene intervals. Counts were converted to transcripts per million (TPM) within each sample (reads per kilobase divided by the per-sample sum, ×10⁶). ORF-level TPM was summed by KO and by functional pathway to yield whole-community functional abundances. Bacterial-community abundance was computed by classifying each contig as archaeal or bacterial from the domain assignment of the eggNOG orthologous groups of its constituent ORFs, taking the majority call. ORFs on archaeal contigs were excluded, and the remaining per-ORF TPM was renormalised against the total bacterial TPM of that sample before seasonal ratios were taken. This removes the inflation of bacterial function ratios caused by the decline in archaeal read fraction from ∼58% in summer to ∼21% in winter.

### Taxonomic assignment of BGC-bearing contigs

Contigs were classified taxonomically with Kaiju v1.7.4 against the nr_euk protein reference database. Protein-coding genes predicted by Prodigal were assigned taxonomic labels, and contig-level taxa were determined by the most frequent assignment among the constituent genes at each rank (domain to species). In case of ties, contigs were conservatively designated as unassigned. To ensure consistency with genome-resolved analysis, NCBI taxon names were mapped to GTDB phyla.

### Whole-community composition and amplicon-metagenome comparison

Metagenomic community composition was estimated by assigning contigs to taxa using Kaiju v1.7.4 and weighting each classification by contig abundance (CoverM trimmed-mean coverage). Relative abundances were summarized at phylum and genus levels. These metagenomic profiles were compared, sample by sample, with genus-level 16S rRNA relative abundances from the same four sediments of our previous study (Ullah et al., under review in *Environmental microbiome*), aggregated to phylum. Metagenomic composition is reported as the proportion of taxonomically classified coverage, with the unclassified coverage stated separately. Because Kaiju resolves NCBI taxonomy and the amplicon data use GTDB taxonomy, genus-level comparisons were restricted to key lineages, with synonyms reconciled explicitly (e.g., GTDB *Franzmannia/Billgrantia* corresponding to NCBI *Halomonas*; GTDB *Moraxella_A* corresponding to NCBI *Moraxella*).

### Genome-resolved biosurfactant gene panel

Because the antiSMASH-based analysis is specific to NRPS lipopeptide clusters, a gene-targeted screen was added spanning the broader biosurfactant and amphiphile range. Each of the 44 MAGs, and separately the whole assembly, was screened for a curated panel covering the lipopeptide, glycolipid, bioemulsifier and membrane-amphiphile classes. Candidate genes were identified from three complementary annotation layers of each predicted ORF including the eggNOG preferred gene name, the KEGG Orthology assignment and the Pfam domain composition, and independently by homology to characterized reference proteins.

Several panel genes share names or sequence homology with unrelated enzymes, so annotation-based presence alone can be misleading; we therefore added a verification step to separate genuine biosurfactant determinants from such false friends, resolving two recurrent cases explicitly. First, rhamnolipid (*rhl*) name-matches were retained only if their Pfam domain composition was consistent with a rhamnosyltransferase, this effectively excludes ATP-dependent RNA helicase RhlB (Pfam DEAD/Helicase_C), which shares the gene symbol. Second, hits to the NRPS-dependent lipopeptide classes (surfactin, iturin, fengycin, lichenysin) were retained only where the coding ORF carried a genuine NRPS module i.e., a condensation and an adenylation (AMP-binding) domain on the same protein. Domain composition for this verification was taken from a per-sequence Pfam scan (hmmsearch against Pfam-A v35.0, per-domain i-value < 1×10⁻⁵), not from the eggNOG inherited from the reference orthologue Pfam field, because the latter reports the domains of the reference orthogroup rather than of the query and produced systematic false positives (e.g., *ppsA*/plipastatin calls that carried no NRPS domain on direct scanning). This step excluded non-NRPS homonyms such as *ppsA* (pyruvate-water dikinase / PEP synthase).

To ensure transparency regarding the impact of this filtering, each biosurfactant class is reported under two distinct criteria: survey presence (any panel gene detected) and the stricter verified criterion (annotation-confirmed determinants only). Finally, every metagenome-assembled genome’s ORF was comprehensively screened for the rhamnosyltransferases RhlB and RhlC using a combination of KEGG Orthology (KO) identifiers, Pfam domain signatures, and gene name queries.

### Proteome-acidity analysis of osmoadaptation strategy

The salt-in versus salt-out osmoadaptation strategy of each recovered genome was assessed annotation-free from proteome acidity, an established genome-wide compositional signature of halophilic adaptation (Oren 2013; Paul et al. 2008) For each MAG, proteome acidity was calculated as the genome-wide percentage of acidic (Asp + Glu) minus basic (Lys + Arg) residues across all predicted proteins. An excess of acidic residues is diagnostic of salt-in strategy in which cytoplasmic proteins function at molar intracellular KCl; near-neutral proteome acidity is characteristic of salt-out strategies.

### Targeted assessment of biosurfactant biosynthesis

The hypothesis of lipopeptide biosurfactant biosynthesis was tested through six independent lines of evidence.

### Manual verification of putative biosurfactant genes

Candidate KO and gene-name hits to known biosurfactant producing genes (*rhl*, *srf*, *fen*, *itu*) were verified manually against KO, Pfam, and eggNOG orthology annotations to exclude false hits arising from non-biosurfactant functions.

### Curated orthology profiling

As an independent test using externally defined thresholds, all predicted proteins were searched with HMMER v3.4 against KOfam profile hidden Markov models for 57 KEGG orthologs covering the core biosurfactant genes of our previous genome-mining panel (Ullah et al. 2026) together with their precursor and accessory genes: phosphopantetheinyl transferases (*sfp*, *acpS*, *entD*), acetyl-CoA carboxylase subunits, fatty-acid synthesis (*fabB*, *fabD*, *fabF*, *fabG*, *fabH*, *fabI*, ACP), dTDP-rhamnose synthesis (*rmlA-D*), trehalose synthesis (*otsA*, *otsB*, *treS*), UDP-sugar supply (*galU*, *galE*), sulfolipid (*sqdB*, *sqdX*), ornithine lipid (*olsA*, *olsB*), and acyl-homoserine-lactone synthases. KEGG’s curated per-family bit-score thresholds were applied, using the sequence-level cutoff (-T) for orthologs with a “full” score type and both sequence- and domain-level cutoffs (-T with --domT) for those with a “domain” score type. Because this method applies externally curated thresholds and imposes no requirement for genes to be co-located, it provides a permissive community-level count independent of assembly contiguity.

### Condensation-domain subtype classification

Condensation domains catalyse peptide-bond formation in NRPS assembly lines and fall into subtypes that reflect their position and function; the Starter subtype initiates synthesis by attaching a fatty acid, and its presence is therefore diagnostic of lipopeptide assembly. Condensation domains within NRPS and NRPS-like regions were extracted from antiSMASH GenBank feature records and classified by the subtype recorded in the domain_subtypes qualifier, which applies the profile models of Rauch et al., (2007). All 20 condensation domains carried a subtype assignment, so the survey is complete rather than partial. Presence of a Starter subtype at the N-terminus of module 1 was used as the diagnostic marker of a lipopeptide-initiating NRPS system.

Adenylation (A) domains were extracted from antiSMASH GenBank files with BioPython v1.79 (23) (AMP-binding and A-OX domains >50 amino acids) and substrate specificity prediction were obtained using the antiSMASH Active Site Finder module (24) (Stachelhaus code).

### Assembly-independent read-level search

Because large NRPS genes assemble poorly, lipopeptide potential was also assessed directly in the unassembled reads. Reads from all four samples were searched with DIAMOND blastx (best hit per read) against the MIBiG v3.1 protein set, assigned to a biosynthetic class by their best-hit cluster, and binned by amino-acid identity. Because MIBiG cluster boundaries include flanking primary-metabolic and tailoring genes, reads assigned to a cluster in this way were re-examined against the full reference set to distinguish matches to synthetase modules from matches to cluster-flanking enzymes. To control for the conserved condensation and adenylation domains shared by all NRPS, the identity distribution of reads assigned to lipopeptide clusters was compared with that of reads assigned to other (non-lipopeptide) NRPS; a genuine lipopeptide gene would yield an excess of high identity reads over this background.

### Reference protein homology screen

Several biosurfactant and amphiphile classes are not recognised by antiSMASH cluster rules, either because they are glycolipids and bioemulsifiers rather than modular assembly lines, or because their pathways are dispersed rather than clustered and are consequently absent from MIBiG. All predicted proteins were therefore screened by DIAMOND blastp against a curated reference of characterized enzymes comprising: 42 MIBiG v3.1 biosurfactant clusters (572 proteins) spanning the lipopeptides (surfactin, iturin, fengycin, lichenysin, arthrofactin, putisolvin, viscosin, viscosinamide, orfamide, entolysin, WLIP, syringomycin, syringopeptin, polymyxin), glycolipids (mannosylerythritol lipid, ustilagic acid) and the emulsan bioemulsifier; rhamnolipid RhlA/RhlB/RhlC; and UniProt reference proteins for classes associated with the lineages recovered here but absent from MIBiG, actinobacterial trehalose mycolates (Pks13, PapA5, mycolyltransferases Ag85A/B/C, FadD32), ornithine lipids (OlsA, OlsB and the bifunctional OlsF), sulfoquinovosyl diacylglycerol (SqdB and the archaeal homologue Agl3) and diacylglyceryl-trimethylhomoserine (BtaA, BtaB).

A protein was counted as a genuine determinant only at ≥60% identity and ≥70% coverage of the reference. A permissive tier (≥40% identity, ≥50% coverage) was evaluated in parallel for classes whose only characterized references derive from distantly related taxa, so that absence is not asserted on stringency alone. Class-specific committed steps were required rather than any panel member: for rhamnolipid, *rhlA* together with the co-occurring *rhlA-rhlB-rhlC* set, since RhlB and RhlC are rhamnosyltransferases shared with lipopolysaccharide O-antigen biosynthesis and an RhlB/RhlC hit lacking *rhlA* was assigned to LPS; for trehalose mycolates, the mycolyltransferases Ag85A/B/C, since Pks13 and PapA5 share ketosynthase and acyltransferase domains with unrelated polyketide systems.

### External validation in an independent genome catalogue

To test the generality of the biosurfactant results, the gene-panel, antiSMASH/MIBiG and reference-protein screens, including the trehalose-mycolate and membrane-amphiphile references, were repeated on an independent catalogue of 1,773 metagenome-assembled genomes from Qaidam Basin desert soils (Liu et al. 2025). Genes were predicted with Prodigal v2.6.3 (-p meta) and screened the same way as as described above.

### Membrane-lipid remodeling and exopolysaccharide analysis

To characterize the community’s surface-active-lipid response, ORFs were assigned to membrane-lipid remodeling and exopolysaccharide (EPS) functions using KO and preferred-name annotations, verified by Pfam domain presence.

Membrane-remodeling functions included phospholipase A and lysophospholipase (lysophospholipid generation), fatty-acid desaturase, and the phosphorus-free membrane lipids ornithine lipid and sulfoquinovosyl diacylglycerol. Desaturase abundance was quantified from the FA_desaturase Pfam domain to exclude mis-annotated homologues, including a TonB-dependent receptor incorrectly labelled *desA*. Ornithine-lipid and sulfolipid abundances were taken from the sequence-verified homology hits rather than from inherited from the reference orthologue gene names, which proved unreliable in both directions for these classes. To test whether phosphorus-free lipid synthesis reflected phosphate limitation, the phosphate-starvation regulon was quantified in parallel (*phoB*/*phoR*; the *pst*/*phoU* high-affinity transport system; alkaline phosphatases *phoA*/*phoD*; phosphonate utilisation *phn*; *phoH*; and polyphosphate kinase/exopolyphosphatase), with a housekeeping set (*rpoB*, *gyrB*, *recA*, *rpsB*, *dnaK*) for normalization. Cyclopropane-fatty-acid and cardiolipin synthases were also quantified but did not show robust winter enrichment within the bacterial community and are not interpreted as seasonal responses.

EPS and capsule-related functions were quantified from CAZy annotations and gated on polysaccharide export systems to distinguish secreted polysaccharide potential from generic envelope glycosylation. Specifically, the *Wza*-*Wzc* capsule export system and group-2 capsular Kps system was quantified, with the *Wzy*-dependent O-antigen route (*Wzx*/*Wzy*) used as a specificity control (Whitfield 2006). All functions were quantified as whole-community TPM, recalculated within the bacterial community, and compared between seasons. Verified genes were attributed to source genomes by contig-to-bin association and DIAMOND self-mapping, and confirmed at KO and Pfam levels and, where applicable, by operon-level contiguity.

### Statistical analysis

Analyses were performed in R v4.4.3. Associations of per-MAG biosynthetic richness with genome size, coding density, and seasonal relative abundance were tested by Spearman rank correlation across the 44 MAGs, and the genome-size effect was summarised as the slope of a linear fit. Biosynthetic richness was compared between seasonally restricted and year-round cohorts by Mann–Whitney U test. Seasonal community structure was tested by PERMANOVA on Bray-Curtis distances of the 16S profiles (13 samples, 999 permutations), with homogeneity of multivariate dispersion assessed by PERMDISP (betadisper) to confirm that a significant PERMANOVA reflects a compositional (location) shift rather than unequal within-season spread; genus-level indicator value (IndVal.g) analysis (indicspecies, 999 permutations) was used to identify statistically significant seasonal indicator taxa. Principal-coordinates ordination was computed in QIIME 2 (Bolyen et al. 2019). To validate the metagenomic community assignments, the seasonal direction of each genus was determined independently from the amplicon data as the sign of the log₂ ratio of mean winter to mean summer genus-level relative abundance and compared with the metagenomic assignment.

### 16S rRNA gene amplicon analysis

Amplicon sequences from the previous 13-sample study were processed as described our previous study (Ullah et al., under review in *Environmental microbiome*). Briefly, reads were denoised into amplicon sequence variants with DADA2 (Callahan et al. 2016) in QIIME 2, taxonomy was assigned against the GTDB reference database, and genus-level relative abundances, Bray-Curtis distances, and principal-coordinates ordination were computed within QIIME 2 for the community-shift and cross-validation analyses reported here.

## Results

### Genome-resolved view of a seasonally structured hypersaline community

Metagenomic sequencing of four sediment samples across summer (C3, C6) and winter (H2, H6) yielded 924.88 million raw paired-end reads (Q30 95.61–97.12%), of which 915.06 million (98.94%) were retained after quality filtering. Independent assembly recovered 2,993,427 contigs ≥500 bp (N50 891–1,053 bp), from which 5,172,059 protein-coding ORFs were predicted (Table 2). Binning and dereplication yielded 44 medium-to-high-quality MAGs distributed across ten bacterial and archaeal phyla. Recovered genomes spanned the ecological coverage expected of hypersaline sediments, including salt-adapted archaea (*Halobacteriota*, *Nanohaloarchaeota*), *Bacteroidota, Pseudomonadota, Chloroflexota,* and *Actinomycetota*.

Ordering genomes by their winter-to-summer abundance ratio resolved three groups (Figure 1A). Summer-favoring MAGs were dominated by salt-in archaea (*Haloarcula, Halobaculum, Haloferax*) and *Salinibacter*, which was the most strongly summer-associated genome recovered. Winter-favoring MAGs comprised salt-out bacteria including *Pseudomonadota*, *Bacteroidota*, and *Myxococcota* (*Moraxella_A, Loktanella, Roseovarius, Gillisia, Persicimonas, and Halomonas_B*). A smaller set showed little seasonal preference.

**Figure 1.**
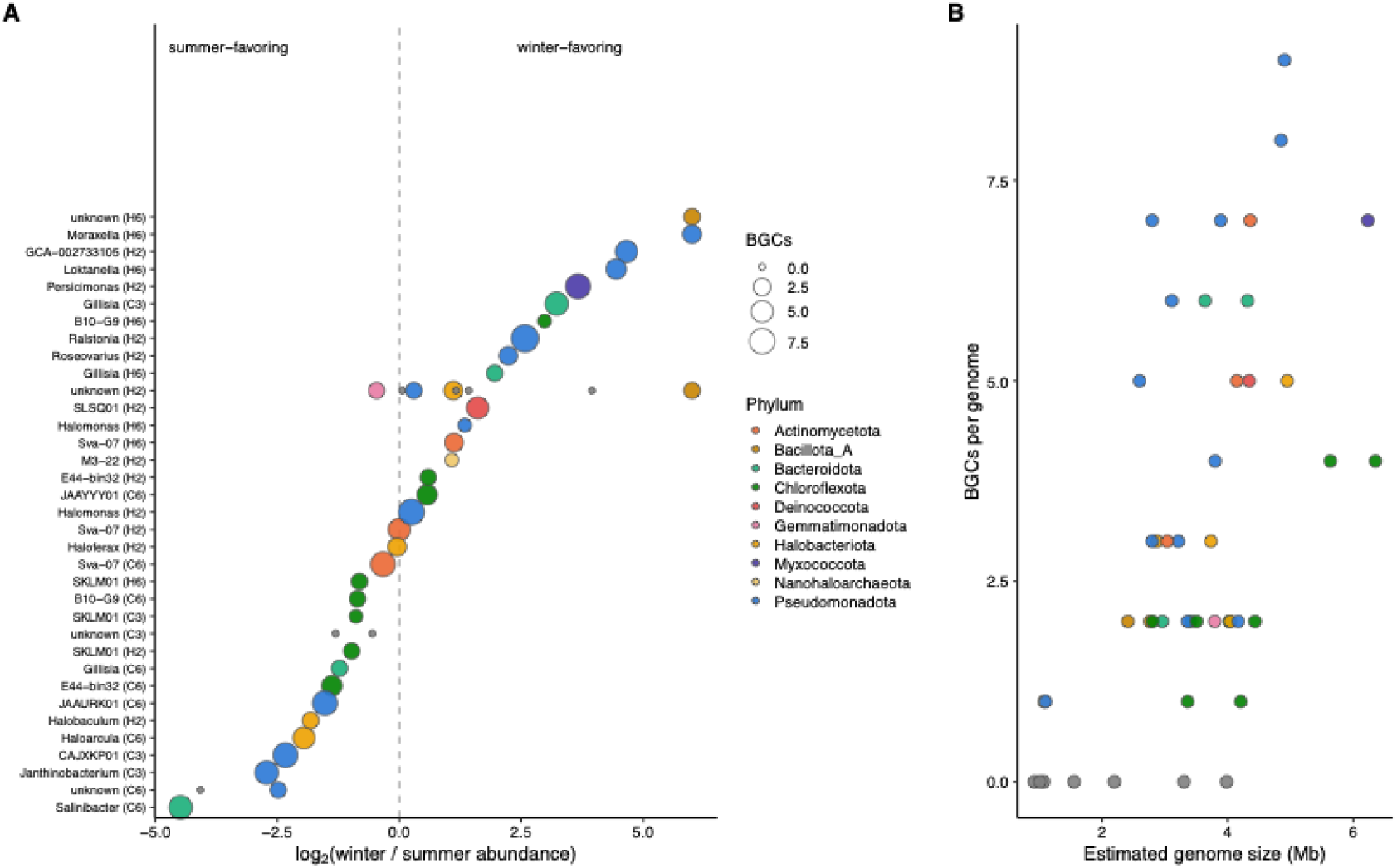
Genome-resolved composition of the Qaidam Basin hypersaline sediment community. Forty-four metagenome-assembled genomes (MAGs) recovered from four sediment samples covering two seasons (summer: C3, C6; winter: H2, H6), coloured by GTDB phylum. (A) MAGs ordered by their seasonal abundance preference, expressed as log₂ of the ratio of mean winter to mean summer trimmed-mean coverage; points left of zero are summer-favoring, right of zero winter-favoring. Point size denotes the number of biosynthetic gene clusters (BGCs) encoded by each genome. Salt-in archaea (Halobacteriota, Nanohaloarchaeota) and Salinibacter dominate the summer-favoring communities, whereas salt-out bacteria dominate the winter-favoring communities. (B) Genome size versus BGC count per genome, showing a positive association between genome size and biosynthetic capacity. Abundance is expressed relative to the whole community, because MAGs represent ∼6% of assembled contigs, seasonal preferences describe the recovered fraction and are validated at the whole-community level in Figures 5-2 and 5-3.

Genome size scaled positively with biosynthetic capacity: larger *Pseudomonadota*, *Myxococcota*, and *Chloroflexota* genomes encoded up to nine BGCs, whereas streamlined salt-in archaea and *Nanohaloarchaeota* carried few or none (Figure 1B). Biosynthetic richness scaled with genome size (Spearman ρ = 0.56, *p < 0.001*) and coding capacity (ρ = 0.63, *P < 0.001*), with no evidence of streamlining in coding density (ρ = −0.06, *p = 0.70*) and no association with seasonal abundance (ρ = 0.18, *p* = 0.24). The archaeal share of mapped reads fell from a summer mean of ∼58% comprising 74% in C3 and 42% in C6 to a winter mean of ∼21% comprising 30% in H2 and 11% in H6, reflecting a seasonal shift.

### 16S rRNA profiling confirms seasonal transition and validates recovered genomes

To place MAGs in whole-community context and test whether they represent the broader community, we compared genome-resolved abundance against 16S rRNA amplicon profiling across the full 13-sample sediment series (Figure 2, Table 3). Community composition differed significantly between seasons (PERMANOVA on Bray-Curtis distances, R² = 0.71, *p* = 0.002), with statistically homogeneous within-season dispersion (PERMDISP *p* = 0.88), indicating a genuine compositional shift rather than unequal within-season variability. Summer communities were enriched in salt-in *Halobacteriota* (*Haloarcula, Halobaculum, Haloferax*) and *Nanohaloarchaeota*, whereas winter communities transitioned toward salt-out bacteria, including *Halomonadaceae* (annotated in GTDB as *Billgrantia* and *Franzmannia* and corresponding to the *Halomonas* lineage recovered as MAG *Halomonas*_B), *Rhodobacteraceae* (*Loktanella, Roseovarius*), *Flavobacteriaceae* (*Gillisia*), and *Moraxellaceae* (*Moraxella*_A) (Figure 2A, 5-2B).

**Figure 2.**
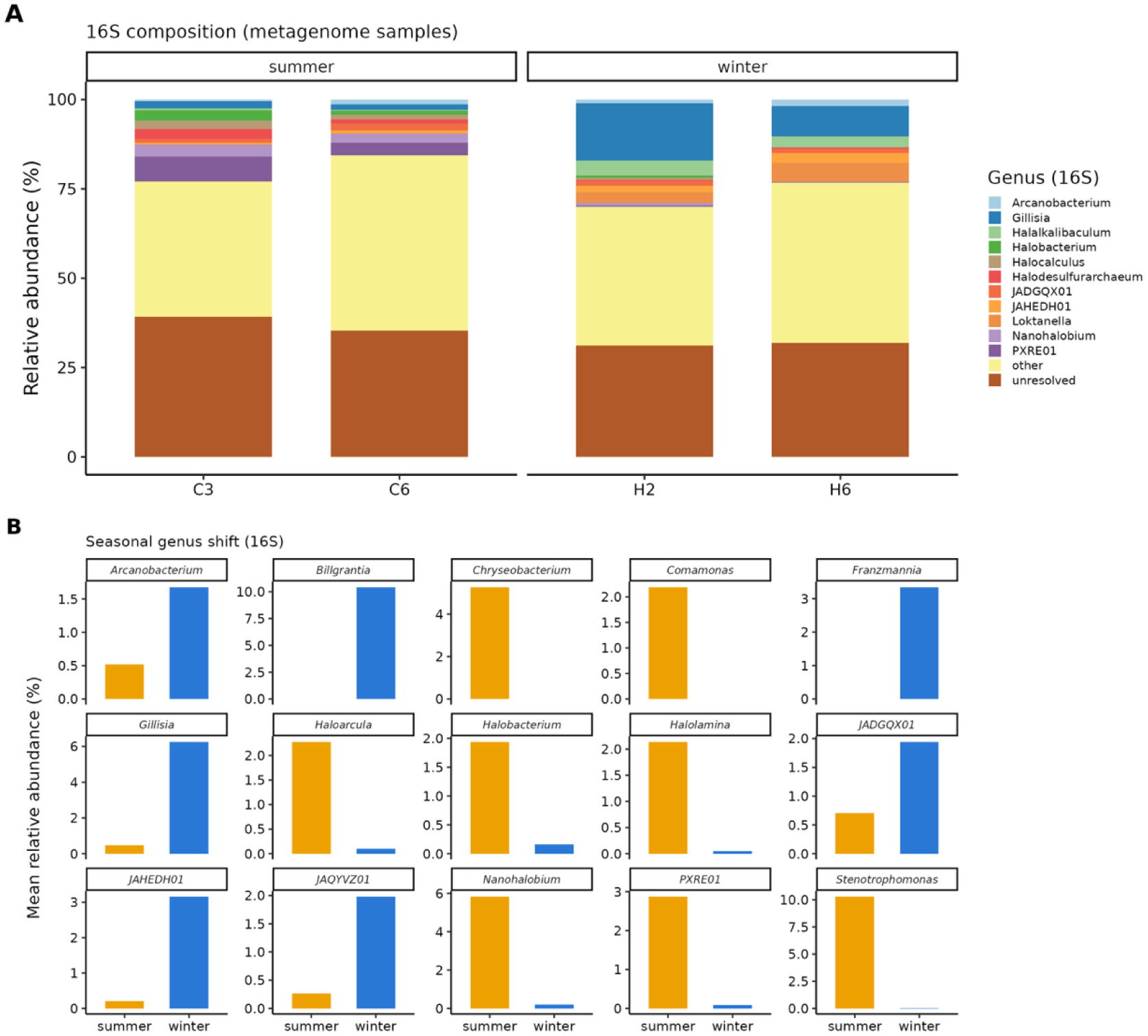
16S rRNA amplicon profiling confirms seasonal community shift across the full sediment series. (A) Genus-level community composition (16S rRNA, GTDB taxonomy) of the four metagenome samples, grouped by season; the twelve most abundant genera are shown, with remaining taxa collapsed into “other” and taxonomically unassigned reads into “unresolved.” (B) Mean relative abundance of abundant genera in summer versus winter across the full sediment series, illustrating the seasonal shift from salt-in archaea to salt-out bacteria. Under the current GTDB taxonomy the Halomonas lineage (recovered as MAG Halomonas_B; Figure 1) is annotated as Billgrantia and Franzmannia; genus names follow GTDB throughout.

**Table 3.** Genome-resolved and 16S rRNA relative abundance of MAG-recovered as genera.

| Genus | Phylum | Metagenom<br>(summer /<br>winter) | 16S rRNA %<br>(summer /<br>winter) | Seasonal<br>preference<br>(MG \| 16S) |
| --- | --- | --- | --- | --- |
| *Gillisia* | *Bacteroidota* | 0.59 / 3.64 | 0.74 / 9.13 | winter \| winter |
| *Halomonas* | *Pseudomonadota* | 0.41 / 0.61 | 0.01 / 5.07 | winter \| winter |
| *Haloarcula* | *Halobacteriota* | 0.57 / 0.15 | 3.60 / 0.15 | summer \| summer |
| *Halobaculum* | *Halobacteriota* | 0.55 / 0.16 | 1.23 / 0.22 | summer \| summer |
| *Loktanella* | *Pseudomonadota* | 0.03 / 0.57 | 0.07 / 2.72 | winter \| winter |
| *Salinibacter* | *Bacteroidota* | 0.40 / 0.02 | 1.12 / 0.04 | summer \| summer |
| *Roseovarius* | *Pseudomonadota* | 0.06 / 0.29 | 0.00 / 0.01 | winter \| winter |
| B10-G9 | *Chloroflexota* | 0.48 / 0.74 | 0.03 / 0.10 | winter \| winter |
| CAJXKP01 | *Pseudomonadota* | 0.17 / 0.03 | 0.11 / 0.01 | summer \| summer |
| GCA-002733105 | *Pseudomonadota* | 0.01 / 0.15 | 0.01 / 0.23 | winter \| winter |
| *Persicimonas* | *Myxococcota* | 0.01 / 0.12 | 0.01 / 0.08 | winter \| winter |
| *Haloferax* | *Halobacteriota* | 0.25 / 0.24 | 0.00 / 0.07 | N.S. \| winter |
| M3-22 | *Nanohaloarchaeota* | 0.12 / 0.25 | 0.47 / 0.00 | winter \| summer |
| E44-bin32 | *Chloroflexota* | 0.22 / 0.16 | 0.01 / 0.29 | N.S. \| winter |

**Note:** NS = Denote genera with no strong metagenomic seasonal difference Table 3 compares MAG
abundance with 16S abundance for the 14 genera for which MAGs were recovered while Figure 2 shows
the 16S seasonal pattern for the most abundant genera. The two sets overlap partially, reflecting
differences in abundance versus recoverability.

Apart from the *Moraxellaceae*, the winter salt-out community is composed of lineages whose described diversity is predominantly marine. *Halomonadaceae*, *Rhodobacteraceae* and *Flavobacteriaceae* are all families in which most characterized genera derive from seawater, coastal or polar marine habitats (Simon et al. 2017; Bowman 2005), and the genera recovered here, *Loktanella, Roseovarius, Gillisia*, are typical members of that marine diversity. The seasonal transition therefore represents not only a switch in osmoadaptation strategy but a shift toward a community of marine-affiliated halophiles, consistent with the physicochemical constraints shared between marine and hypersaline environments.

Across the 14 genera recovered as MAGs, metagenomic abundance was positively rank-correlated with 16S relative abundance (Spearman ρ = 0.54), and seasonal preference was consistent between methods for 11 of 14 genera (Table 3). Alignment was complete among ecologically dominant taxa: the winter salt-out genera *Halomonas, Gillisia, Loktanella*, and *Roseovarius*, and the summer salt-in genera *Haloarcula, Halobaculum*, and *Salinibacter* were assigned the same seasonal preference by both approaches.

Beyond the recovered genomes, the entire metagenomic community was classified independently with Kaiju, weighted by contig coverage, and compared it sample wise with the 16S composition (Table 4). At phylum level, the salt-in archaea *Halobacteriota* dominated summer sediments on both approaches (16S 52%, metagenome 44% of classified coverage) and collapsed in winter (5% and 13%), whereas the salt-out bacteria *Pseudomonadota* and *Bacteroidota* rose in winter on both (*Pseudomonadota* 37%/36%; *Bacteroidota* 26%/30% in 16S/metagenome). At genus level, 10 of 12 ecologically defining lineages agreed on seasonal direction between these two methods: the salt-in haloarchaea (*Haloarcula*, *Halobaculum*, *Halonotius*, *Halolamina*) and the salt-in bacterium *Salinibacter* were summer-associated on both, and the salt-out bacteria, the *Halomonas* lineage (GTDB *Franzmannia*/*Billgrantia*), *Moraxella*, *Loktanella*, *Gillisia* and *Persicimonas*, winter-associated on both. The two exceptions (*Haloferax*, *Roseovarius*) were near the amplicon detection limit (<0.1% of 16S reads), and the metagenome placed each in its biologically expected season (*Haloferax*, a haloarchaeon, in summer; *Roseovarius*, a *Rhodobacteraceae*, in winter). Between 52% and 65% of metagenomic coverage derived from contigs that could not be resolved to phylum against nr_euk. This was consistent with a community rich in novel halophilic lineages; DPANN *Nanohaloarchaeota*, detected by 16S in summer (∼6%), were similarly under-recovered by reference-based classification (Narasingarao et al. 2012).

**Table 4.**
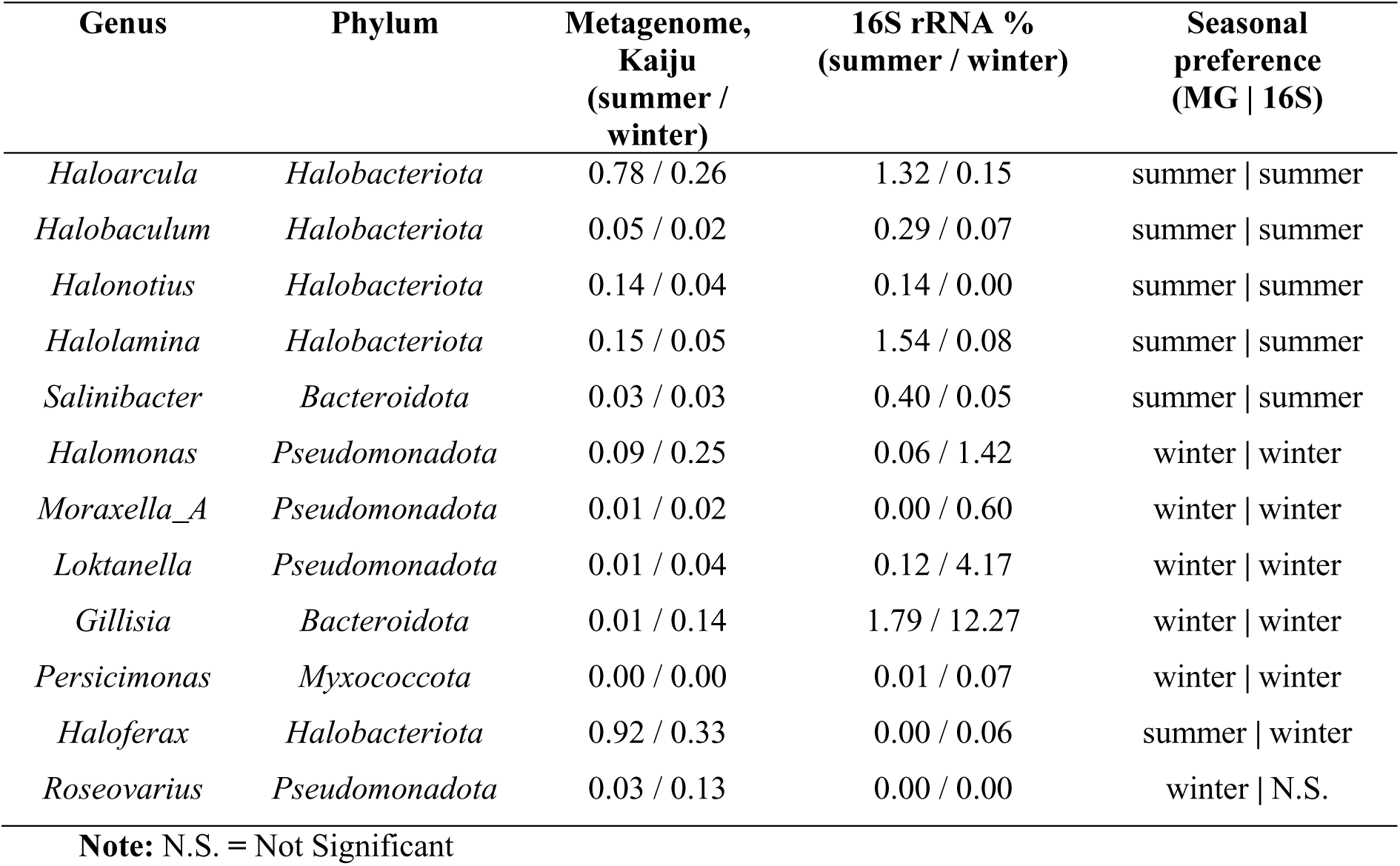
Amplicon vs metagenomic community composition.

### A seasonal salt-in to salt-out osmoadaptation switch

The seasonal restructuring of the community corresponds to a switch between salt-in and salt-out strategies for cytoplasmic osmotic balance (Figure 3). The recovered archaea (*Haloarcula, Halobaculum, Haloferax*) formed a distinct acidic cluster (acidic minus basic > 5.5%), consistent with salt-in proteomes, whereas salt-out bacteria clustered near neutrality (1–4%). *Salinibacter*, the known salt-in bacterium, occupied the expected intermediate position, providing an internal control. Acidic salt-in archaea were summer-favoring, and near-neutral salt-out bacteria were winter-favoring, directly coupling proteome acidity to the seasonal abundance axis (Figure 3A).

**Figure 3.**
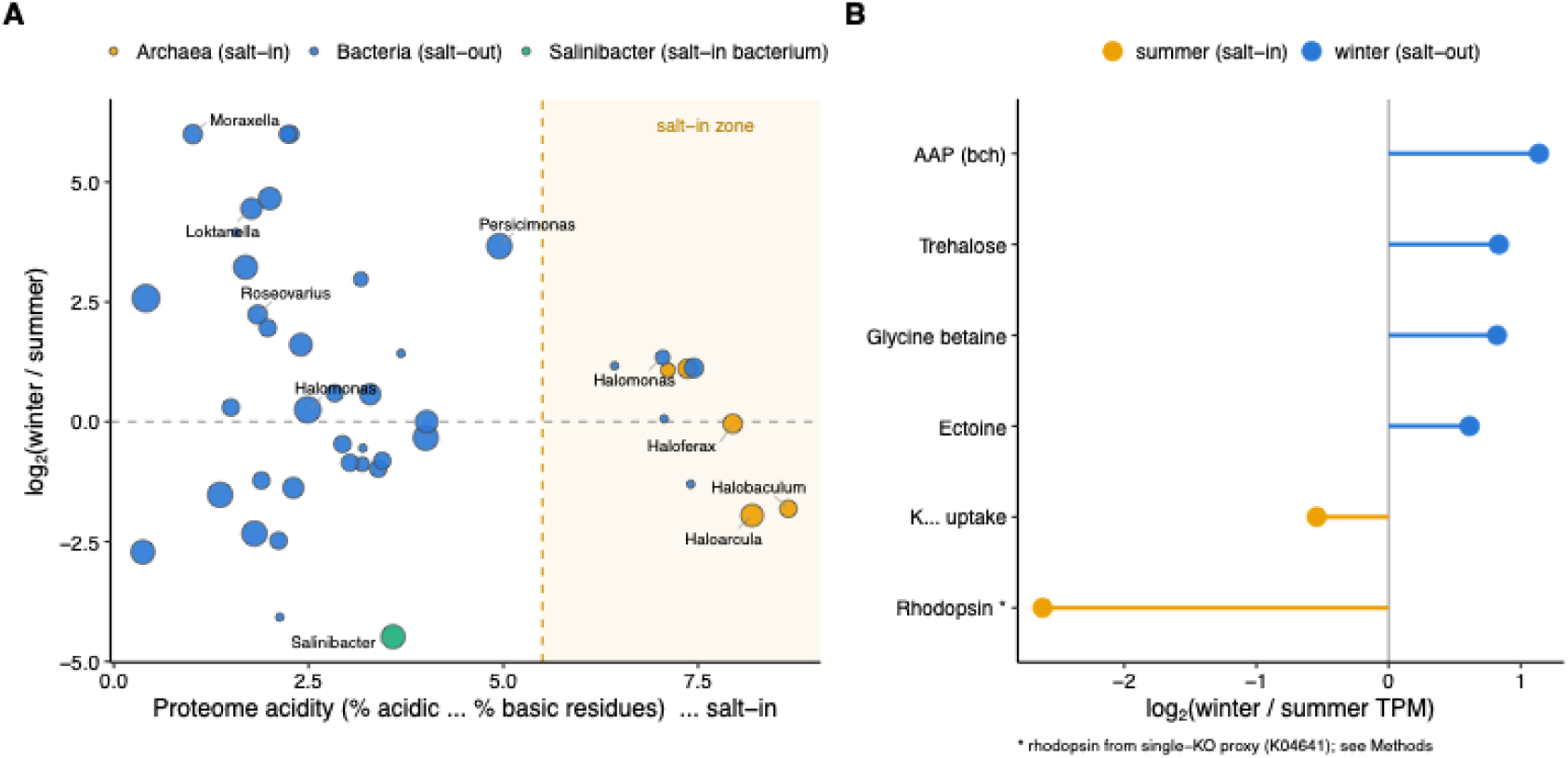
A seasonal salt-in to salt-out osmoadaptation switch**. (A)** Proteome acidity of each MAG, computed annotation-free as the genome-wide percentage of acidic (Asp + Glu) minus basic (Lys + Arg) residues (x-axis; higher values indicate salt-in adaptation), plotted against seasonal abundance preference (y-axis, as in Figure 1A). Points are coloured by osmoadaptation community; point size denotes BGC count. The shaded region marks the salt-in zone; salt-in archaea occupy it and are summer-favoring, salt-out bacteria cluster near neutrality and are winter-favoring, and the salt-in bacterium *Salinibacter* occupies the expected intermediate position (internal control). **(B)** Whole-community functional abundance of osmoadaptation-associated pathways, expressed as log₂ of the winter-to-summer ratio of summed TPM; compatible-solute biosynthesis is winter-enriched (salt-out) while rhodopsin phototrophy is summer-enriched (salt-in). Potassium-uptake abundance is shown for completeness but not interpreted directionally, as these systems occur in both communities. Asterisk: rhodopsin abundance is based on a single conservative KO marker (K04641) and is likely an underestimate of the true summer signal. Values are whole-community TPM ratios; bacterial-community values are given in the text and are lower for trehalose (1.17-fold) and glycine betaine (1.29-fold).

Whole-community functional abundance validated this osmoadaptation switch (Figure 3B). Compatible-solute biosynthesis pathways (ectoine, trehalose, glycine betaine), characteristic of salt-out strategies, were winter-enriched and retained their winter direction after restricting the analysis to the bacterial fraction. Within the bacterial community, ectoine biosynthesis (ectA/B/C/D) was ∼1.5-fold higher in winter than summer, while trehalose (∼1.2-fold) and glycine-betaine (∼1.3-fold) synthesis showed more modest winter biases and are treated as supporting signals. On the salt-in side, the summer archaea had acidic proteomes, dominated the summer read pool, and carried microbial rhodopsins, which were summer-enriched at the community level (winter/summer ratio ∼0.39).

### Surface-active-lipid response: membrane remodeling and capsule export

We next asked how the community’s potential in surface-active amphiphilic lipids is structured seasonally (Figure 4). Two coordinated responses were found, both winter-enriched and both localised to salt-out bacteria.

**Figure 4.**
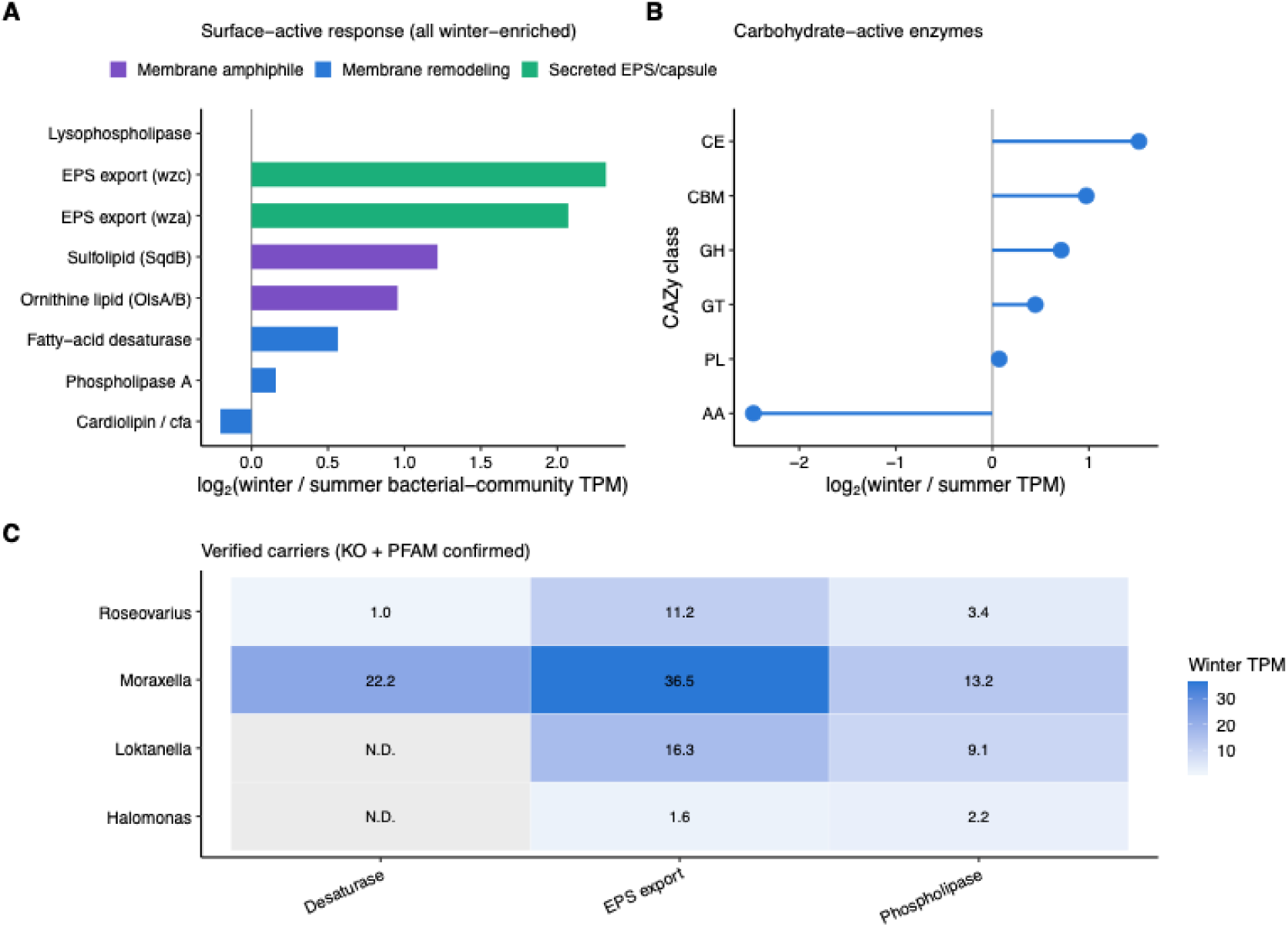
The community’s surface-active-lipid response is winter-enriched membrane remodeling and secreted exopolysaccharide. (A) Log2 fold-change (winter/summer) in community-level transcript abundance (TPM) for surface-active genes, grouped by functional category (membrane amphiphile, membrane remodeling, secreted EPS/capsule). (B) Log2 fold-change (winter/summer) in TPM for CAZyme classes (CE, CBM, GH, GT, PL, AA). (C) Heatmap of winter TPM for KO- and PFAM-confirmed carriers of decarboxylase, EPS export, and phospholipase genes across four genera (Roseovarius, Moraxella, Loktanella, Halomonas); N.D. = not detected.

The first response is membrane-lipid remodeling, comprising both compositional adjustment and lipid substitution. Fatty-acid desaturase showed the strongest and most consistent single membrane signal, with ∼2.2-fold winter enrichment within the bacterial fraction (Figure 4A).

Phospholipase A was also winter-enriched (∼1.5-fold within bacteria); lysophospholipase abundance was only weakly elevated (∼1.1-fold) and is treated as a supporting signal. Alongside this transition, the community shifted toward phosphorus-free membrane lipids: ornithine-lipid synthesis (OlsA/OlsB) was enriched ∼1.9-fold within the bacterial community, confirmed independently by curated KOfam profiles (OlsB 4.7-fold, OlsA 2.7-fold winter) (Figure 4A). Sulfoquinovosyl-lipid capacity was incomplete: the precursor synthase SqdB was detected across all samples (6–49 hits, without seasonal enrichment) but the core sulfoquinovosyltransferase SqdX was essentially absent (0–1 hits), indicating precursor capacity without complete sulfolipid synthesis. Ornithine-lipid genes were carried by *Alphaproteobacteria*, principally *Rhizobiales*-affiliated contigs, and were not associated with archaeal contigs. The bifunctional ornithine-lipid route (OlsF) was not detected, indicating synthesis through the two-step OlsB-OlsA pathway, while diacylglyceryl-trimethylhomoserine was detected on too few ORFs to interpret. This shift was not accompanied by a phosphate-starvation response: no arm of the Pho regulon exceeded 1.4-fold within the bacterial community, and high-affinity phosphate transport (*pst*/*phoU*) and phosphonate scavenging (*phn*) were not winter-enriched (∼0.8–fold each), against a housekeeping control of 1.0-fold. Cardiolipin and cyclopropane-fatty-acid synthases did not remain winter-enriched after restricting the analysis to bacteria (∼0.96-fold).

The second response is secreted exopolysaccharide (EPS) and capsule export. The *Wza*-*Wzc* capsule export pathways remained winter-enriched within the bacterial community (*wza* ∼2.6-fold, *wzc* ∼2.0-fold, with accessory wzb also elevated), whereas the *Wzy*-dependent O-antigen route (*wzx*/*wzy*) showed no winter enrichment (∼0.4-fold within bacteria). The alternative group-2 (Kps) ABC exporter route was ∼1.0-fold within bacteria and is treated as supporting only. Winter-enriched carbohydrate-active enzyme families within the bacterial community (Figure 4B) were concentrated in storage- and osmolyte-glycan metabolism: maltose/trehalose phosphorylase (GH65, 2.1-fold), α-glucosidases (GH97 2.6-fold, GH31 1.4-fold), glycogen synthase (GT5, 1.7-fold), and the starch-binding module CBM48 (1.2-fold) alongside α-amylase GH13 (1.2-fold), the most abundant family in the dataset. Families acting on structural polysaccharides moved in the opposite direction: cellulose- and cellobiose-active families (GH5, GH9, GH94) and chitinase GH18 were all summer-directed (0.6–0.7-fold). The α-amylase and α-glucosidase families GH13 and GH31 were only weakly enriched within bacteria (1.2- and 1.4-fold) despite apparent 2.0- and 2.2-fold enrichment at the whole-community level, illustrating the compositional inflation described above.

These surface-active functions were carried by winter salt-out bacteria. Among genome-resolved carriers (verified at KO and Pfam levels), *Moraxella* encoded a complete, contiguous *Wza*-*Wzb*-*Wzc* export operon together with phospholipase A and a fatty-acid desaturase, Loktanella carried group-2 capsule export (KpsE-KpsT) plus lysophospholipase, and *Roseovarius* carried a similar complement but was not a significant seasonal indicator (Figure 4C).

### Biosynthetic repertoire and targeted search for lipopeptide biosurfactants

The community encoded an extensive secondary-metabolite repertoire, 1,468 BGCs across the four assemblies, but its composition and a targeted search claim against established lipopeptide biosurfactant biosynthesis as a major community trait (Figure 5). The BGC pool was dominated by terpene and RiPP-like clusters, with type-III PKS, arylpolyene, and RRE-containing clusters also prominent (Figure 5A). NRPS and NRPS-like clusters formed a minority. Most BGCs (∼91% overall and ∼84% of NRPS-class clusters) resided on unbinned contigs (Figure 5B).

**Figure 5.**
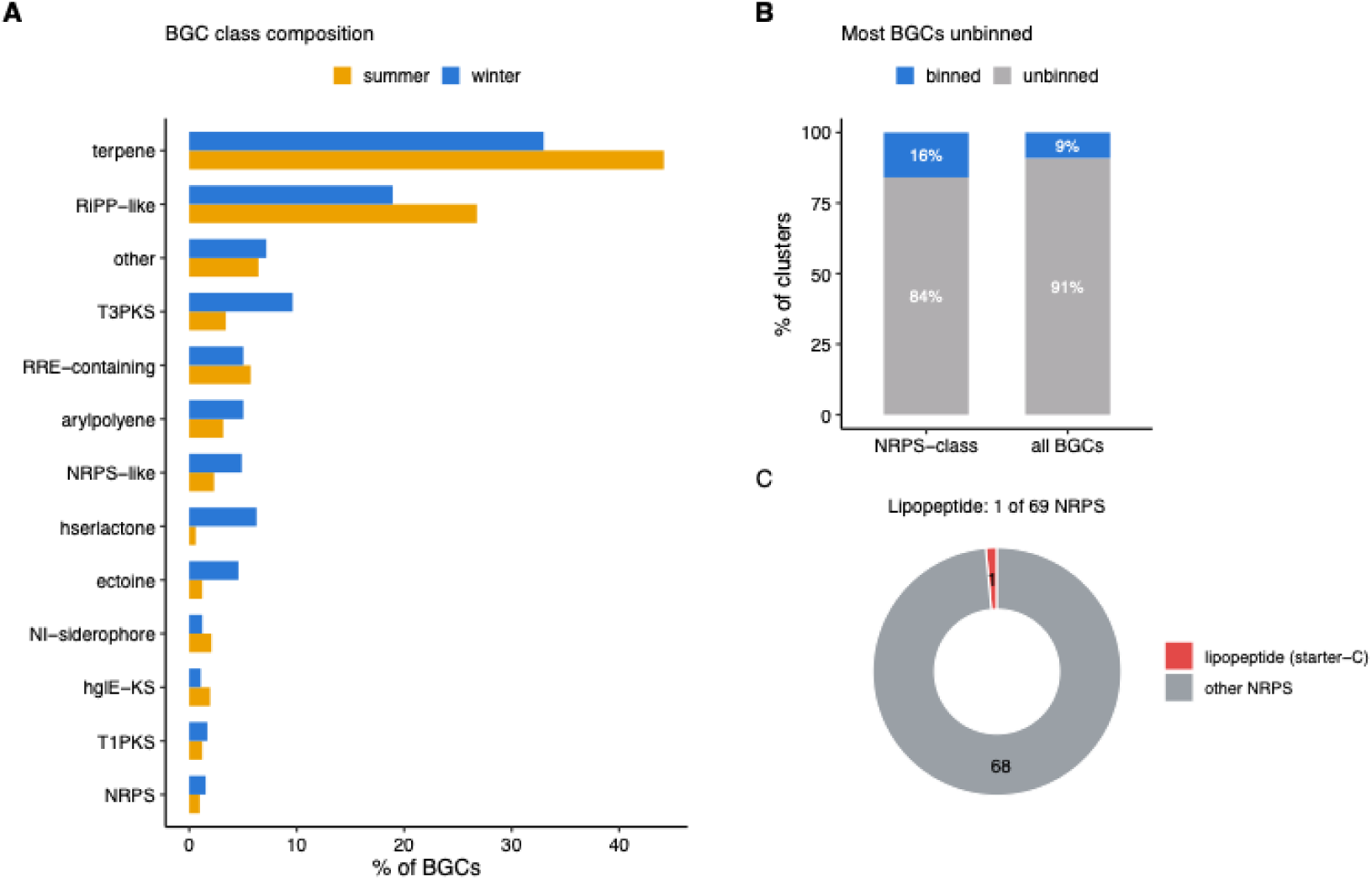
An extensive biosynthetic repertoire in which dedicated lipopeptide biosurfactants are essentially absent. Note: (A) Composition of the 1,468 detected BGCs by product class and season, expressed as the percentage of BGCs in each season (B) Percentage of clusters residing on binned versus unbinned contigs, for all BGCs and for NRPS-class BGCs; the majority of both, and of the NRPS signal specifically, are unbinned. (C) Among all NRPS clusters, the number carrying the Starter condensation domain diagnostic of lipopeptide assembly versus those with standard elongation domain; a single cluster carries the lipopeptide-initiating domain.

Classifying the 1,468 BGC-bearing contigs with Kaiju, independently of the recovered genomes, resolved 41% to phyla distributed across *Pseudomonadota* (231 contigs), *Halobacteriota* (192), *Actinomycetota* (80), *Bacteroidota* (70) and *Bacillota* (14) while 59% resided on contigs of lineages unresolvable against reference databases. Weighting each BGC contig by its coverage showed that this biosynthetic potential follows the seasonal community switch. In summer, the classified biosynthetic signal was dominated by the salt-in *Halobacteriota* (57% of classified BGC coverage), principally *Haloarcula* (27%), with a further 12% from *Actinomycetota*. In winter it shifted to the salt-out bacteria, *Pseudomonadota* (40%) and *Bacteroidota* (37%); the winter *Bacteroidota* signal derived from *Flavobacteriaceae*-affiliated genera (NCBI *Gelidibacter*, *Chitinophaga*, *Polaribacter*, the community recovered as *Gillisia* under GTDB) and the *Pseudomonadota* signal from *Pseudomonas*, *Shewanella* and *Marinobacter*. Between 52% (winter) and 68% (summer) of BGC-bearing coverage derived from unclassifiable contigs, confirming that the community’s biosynthetic potential lies overwhelmingly in novel, unrecovered lineages.

Comparing all detected clusters to characterized references, the dominant matches were compatible-solute biosynthesis (ectoine, 134 matches), followed by pigments (carotenoid 31, flexirubin 20) and siderophores (∼24 combined). Matches to surface-active compound classes numbered seven, all resolved to two-protein hits against fatty-acid-supply genes rather than synthetases. The community’s characterized biosynthetic capacity is therefore directed principally toward osmolyte production and iron acquisition.

Initial KEGG-orthology matches to established biosurfactant genes resolved to false annotations (e.g., *rhlA*/*phaG* and *rhl*-like hits to RNA helicases) rather than genuine biosurfactant biosynthesis. Examining condensation domains directly, of the 20 condensation domains annotated within NRPS and NRPS-like clusters, a single Condensation_Starter domain was recovered (13 LCL, 4 DCL, 2 Dual C/E), residing in one cluster of 69 (Figure 5C).

The single lipopeptide-initiating cluster (sample C6, contig k141_686258, binned in an *Actinomycetota* MAG of the genus *Sva-07*) encoded a starter condensation domain at the N-terminus of module 1 followed by adenylation, carrier-protein, LCL-condensation and a second adenylation domain, the recognized lipoinitiation structure. The cluster was truncated at the contig boundary, no additional condensation domain was present elsewhere in that genome, and KnownClusterBlast returned no match to any characterized MIBiG cluster, indicating an uncharacterized lipopeptide-type system. The single starter domain fell outside the clade formed by the characterized surfactin, fengycin and iturin starter domains. Direct comparison of every NRPS region to experimentally characterized clusters (antiSMASH KnownClusterBlast against MIBiG) returned no match to any validated lipopeptide biosurfactant: the few clusters yielding a lipopeptide name shared only ubiquitous fatty-acid-supply genes (short-chain acyl-CoA synthetase, acetyl-CoA carboxylase) rather than the multi-modular synthetases.

Since large NRPS genes assemble poorly, lipopeptide potential was also assessed directly in unassembled reads. Across the identity distribution, reads assigned to lipopeptide clusters carried the lowest proportion of high-identity hits of any NRPS class, showing no excess over the conserved-domain background shared by all NRPS. The search however retained sensitivity to genuine synthetase sequence. Of the reads assigned to the entolysin cluster, the great majority matched cluster-flanking primary-metabolic and tailoring enzymes annotated within the MIBiG, principally 4-hydroxyphenylpyruvate dioxygenase, rather than synthetase modules. Only two reads, both from C3, matched the entolysin NRPS genes (PSEEN3044, PSEEN3045), at 94–98% amino-acid identity over 49–50 residues, a frequency of approximately one in 10⁸ reads. This scarcity is not explained by absence of the producing lineage as Kaiju assigned 7,612 of 1,248,890 classified C3 contigs (0.61%) to Pseudomonas, the genus most closely associated with lipopeptide production. The candidate lineage is therefore present while its lipopeptide pathway is absent.

An independent search using curated KOfam profiles gave the same result under the KEGG thresholds. Across the fourteen lipopeptide synthetase orthologs of the surfactin, iturin, fengycin and arthrofactin pathways, only two isolated hits were recovered from any sample, one surfactin synthetase C and one plipastatin synthetase E, both in the same winter assembly and no pathway was complete in any assembly. Critically, this search imposed no requirement for genes to be co-located, so the absence is not a product of assembly fragmentation. By contrast, the *Wza*-*Wzb*-*Wzc* capsule export system was detected in all four assemblies and was most abundant in winter (*wza* 38–135 hits per sample), while the *Wzy*-dependent O-antigen route was undetected, reproducing by an orthogonal method the pattern obtained from cluster- and homology-based analyses.

Precursor and accessory genes were quantified alongside the core enzymes. Fatty-acid supply was present throughout (*fabG* 87–519 hits per sample, *fabH* 77–207, *fabI* 31–105, acyl carrier protein 151–299, acetyl-CoA carboxylase subunits 14–112), as was the complete dTDP-rhamnose pathway (*rmlA-D*, 9–123 hits), trehalose synthesis (*otsA* 11–32, *otsB* 26–79) and UDP-sugar supply (*galU* 21–31, *galE* 68–160). Phosphopantetheinyl transferases, which are obligatory for the activation of any NRPS carrier protein, were abundant (*sfp*-type 20–43 hits, AcpS 51–91). The community therefore possesses the precursor supply and the activating pathway required for lipopeptide and glycolipid synthesis with 69 NRPS clusters, while lacking the main synthetases and transferases themselves.

Tracing every hit from ORF to contig to genome bin, 201 of 2,574 hits (8%) resided in metagenome-assembled genomes. Three MAGs carried a core biosurfactant gene. In *Halomonas_B*, an NRPS matching lipopeptide synthetase profiles was resolved by KnownClusterBlast as a siderophore cluster, with all eight significant matches to characterized amphiphilic siderophores (variobactin, potashchelin, pacifibactin, taiwachelin, variochelin, vacidobactin, crochelin, plantaribactin), consistent with its independent antiSMASH classification as an NRP-metallophore. A *Chloroflexota* MAG carried a single *rhlC*-family gene within an O-antigen cluster. The third, the *Actinomycetota Sva-07* MAG described above, carried the only lipopeptide-initiating NRPS in the dataset.

To test whether biosurfactant potential extends beyond the lipopeptide clusters examined above, the 18-gene panel of our genome-mining study was applied to the 44 MAGs. Biosurfactant-associated genes were widespread, 40 of 44 MAGs (91%) carried at least one panel gene, matching the >90% of amplicon reads assigned to putative-producer lineages (Ullah et al., under review in *Environmental microbiome*). However, this signal was carried by near-universal, non-specific genes: fatty-acid synthesis (*fab*, 34 MAGs), alcohol dehydrogenase (*adh*, 23), dTDP-rhamnose synthesis (*rml*, 8) and alginate genes (*alg*, 18). The single biosurfactant class genuinely encoded beyond such housekeeping precursors was the capsular exopolysaccharide export (*Wza*-*Wzc*), present in the salt-out bacteria *Moraxella_A*, *Halomonas_B*, *Gillisia* and *Salinibacter* with the same secreted-capsule/exopolysaccharide presence identified above as the community’s surface-active response.

By contrast, the low-molecular-weight secreted classes were not detected as encoded pathways. Rhamnolipid biosynthesis was not encoded. Gene-name matching returned three *rhlB* hits that direct Pfam scanning identified as ATP-dependent RNA helicases, and the only rhamnose-active enzymes were catabolic rhamnosidases. An independent search using curated KOfam profiles recovered two *rhlA*-family, two *rhlB*-family and three *rhlC*-family sequences across the four assemblies. Comparison against curated references assigned one *rhlA* candidate unambiguously to the PhaG family ((R)-3-hydroxydecanoyl-ACP:CoA transacylase; 52.3% identity to PhaG versus 38.2% to RhlA); the second was equidistant from both families and could not be assigned. Finally, the seven hits occupied seven separate contigs spanning two phyla, no *rhlA* candidate had a co-encoded glycosyltransferase, and the *rhlC*-family hit lay within an O-antigen biosynthesis cluster containing dTDP-sugar pathway genes, polysaccharide synthases and *epsG*, identifying it as a cell-surface rhamnosyltransferase. Since mono-rhamnolipid synthesis requires the co-transcribed *rhlAB* pair, the *rml* and *fab* hits represent sugar and fatty-acid precursors rather than rhamnolipid capacity.

Mannosylerythritol-lipid genes and a dedicated emulsan locus were also absent at the identity and coverage expected of a true orthologue. Actinobacterial trehalose mycolates, the glycolipid class most probable for the *Actinomycetota* present in this community, were also absent: the main mycolyltransferases Ag85A/B/C returned no hit at either stringency, and the only matches to Pks13 and PapA5 covered 14% and 9% of their references, consistent with shared ketosynthase and acyltransferase domains rather than an intact pathway.

The lipopeptide core (surfactin, iturin, fengycin, lichenysin) was absent from all 44 MAGs after verification: 24 MAGs carrying an apparent fengycin/plipastatin hit encoded *ppsA* mis-annotations of pyruvate-water dikinase rather than plipastatin synthetase, consistent with the whole-community finding of a single lipopeptide-initiating cluster among 1,468 BGCs and with the core’s global rarity (0.23% of genomes).

Consistent with these findings, an independent catalogue of 1,773 MAGs from Qaidam Basin desert soils (Liu et al. 2025) also encoded no intact lipopeptide, rhamnolipid, or other dedicated glycolipid biosurfactant pathway, and no trehalose-mycolate pathway (mycolyltransferases, Pks13 and PapA5 were absent from all 1,773 genomes at both stringencies). The membrane amphiphiles enriched in winter here were uncommon in that catalogue, sulfoquinovosyl-lipid synthesis in 47 of 1,773 genomes (2.7%), ornithine-lipid synthesis in 13 (0.7%).

## Discussion

Microbial adaptation to seasonally extreme hypersaline sediments in this environment is built not on freely secreted biosurfactants, but on cell-associated amphiphilic potential: membrane-lipid remodeling and capsular/exopolysaccharide stabilization carried by a winter salt-out community. These cell surface-active adaptations are constrained and enabled by a seasonal transition in osmotic strategy, from summer salt-in archaea to winter salt-out bacteria, which determines which lineages dominate and therefore which amphiphilic responses are even available to be deployed. The surface-active capacity encoded by this community stays attached to the cell rather than being released, and the capabilities present are by the lineages that persist under the prevailing seasonal osmotic conditions.

The community alternates between two coherent seasonal states, organized by osmotic strategy rather than salinity alone. Summer is dominated by salt-in archaea with streamlined genomes and constitutively acidic proteomes adapted to molar intracellular KCl, consistent with the classical haloarchaeal salt-in strategy (Gunde-Cimerman et al. 2018; Oren 2010). These lineages accumulate K⁺ and harvest light through rhodopsins, investing relatively little in secondary metabolism or surface-active lipids. Winter, in contrast, is built on salt-out bacteria that maintain conventional cytoplasmic ionic strength, synthesize compatible solutes such as ectoine, glycine betaine, and trehalose, and deploy cell-surface amphiphilic adaptations instead. The salt-in/salt-out dichotomy emerged independently from four largely non-overlapping signals: the seasonal shift in 16S-based composition across 13 samples, genome-resolved abundance, a proteome-acidity signature that places the known salt-in bacterium *Salinibacter* at its expected intermediate position, and the collapse in the archaeal read fraction from ∼58% in summer to ∼21% in winter. Because these four signals are deemed to be independent evidence, no single methodological error could produce all of them, and the seasonal contrast can be treated as a real property of the community rather than a sampling artefact.

Reading osmotic strategy as the structural constraint also resolves an apparent inconsistency: winter is the more saline season, yet it is dominated by salt-out bacteria, even though salt-out is energetically disfavored at the highest salinities (Oren 2011). The explanation is that the two strategies differ not only in energetic cost but in the biochemical flexibility they permit: the salt-in strategy commits the entire proteome to function at high ionic strength and restricts alternative biochemical responses, whereas the salt-out strategy preserves the capacity to mount coordinated responses to combined stresses (Gutiérrez-Preciado et al. 2024). Winter imposes higher salinity together with cold and desiccation; it is this combination, not salinity alone, that selects salt-out lineages capable of membrane remodeling and capsule production. Osmotic strategy is therefore not simply a response to salt concentration; it determines which further adaptations a lineage can deploy at all, including the surface-active responses.

Within the winter salt-out community, the surface-active response is internally coherent and functionally specific. Membrane remodeling is dominated by fatty-acid desaturase, the strongest and most consistent single seasonal signal in the dataset, enriched ∼2.2-fold in winter within the bacterial community after compositional correction, the expected signature of homeoviscous adaptation, in which increased unsaturation preserves membrane fluidity at low temperature (Chintalapati et al. 2004; Sinensky 1974). Phospholipase A is also winter-enriched (∼1.5-fold within bacteria), consistent with deacylation of membrane phospholipids toward lysophospholipids; lysophospholipase itself is only weakly enriched (∼1.1-fold) and is treated as a supporting signal only. The metabolomics pool detected in the linked amplicon study (Ullah et al., under review in *Environmental microbiome*) is dominated by lysophospholipids, providing independent support for this remodeling pathway. The winter response extends beyond compositional adjustment to lipid substitution: ornithine lipids, phosphorus-free membrane lipids carried by Alphaproteobacterial lineages, are enriched approximately twofold within the bacterial community. The parallel sulfoquinovosyl-lipid route is incomplete, the precursor synthase SqdB is present in all samples without seasonal enrichment, while the transferase SqdX is essentially absent, so sulfolipid substitution is not supported. Substitution of phospholipids by phosphorus-free lipids is a recognised bacterial response to phosphorus limitation (Geiger et al. 2010), but that explanation does not apply here: no arm of the phosphate-starvation regulon is winter-induced, and high-affinity phosphate transport and phosphonate scavenging are, if anything, reduced. The substitution therefore appears to belong to a general envelope-remodeling system rather than to a phosphorus-sparing response. (Cardiolipin and cyclopropane-fatty-acid synthesis, which appear winter-elevated at the whole-community level, lose significance after restricting the analysis to the bacterial fraction, an artefact of the seasonal archaeal decline rather than a genuine bacterial signal, and are not interpreted further.)

Capsule and exopolysaccharide production show comparable specificity. Because envelope glycosyltransferases are abundant and functionally heterogeneous, a winter increase in glycosyltransferase gene counts alone would be uninformative; the EPS interpretation is therefore anchored on export systems rather than on biosynthetic gene families (Whitfield 2006). The *Wza*-*Wzc* capsule export system is winter-enriched within the bacterial community (*wza* ∼2.6×, *wzc* ∼2.0×, with accessory wzb only marginally elevated), whereas the *Wzy*-dependent O-antigen route shows no winter enrichment (∼0.4× within bacteria), which confines the seasonal signal to secreted capsules and EPS rather than to generic cell-envelope glycosylation. The alternative group-2 (Kps) export route does not survive compositional control and is excluded from this interpretation. Winter-enriched carbohydrate-active enzyme families, maltose/trehalose phosphorylase (GH65), α-glucosidases (GH97, GH31) and glycogen synthase (GT5), together with the opposing summer enrichment of cellulose- and chitin-active families (GH5, GH9, GH94, GH18), further support a shift toward osmolyte- and storage-associated glycan metabolism rather than structural polysaccharide degradation. Critically, the same recovered lineages carry both responses: *Moraxella*_A encodes a contiguous *Wza*-*Wzb*-*Wzc* operon alongside phospholipase A and a fatty-acid desaturase, and *Loktanella* carries capsule export plus lysophospholipase. These functions are confirmed at the KO and Pfam level and are significant winter indicators in the independent 13-sample amplicon dataset (Ullah et al., under review in *Environmental microbiome*). Their co-occurrence within the same genomes, combined with community-level enrichment across winter bacterial lineages, indicates that membrane remodeling and capsule production are components of one coordinated, winter-associated envelope strategy of the salt-out community, rather than idiosyncratic traits of a few unrelated taxa.

Despite encoding 1,468 biosynthetic gene clusters across the four assemblies, the community is essentially lacking in lipopeptide biosurfactant production. Among 69 NRPS clusters, only one carries the Starter condensation domain diagnostic of lipopeptide assembly; the remainder encode standard elongation domains characteristic of non-lipopeptide NRPS systems. The deficit extends to glycolipids: no genome or contig encoded a verified rhamnolipid-committed RhlA, the recovered *rhlA*-family candidates were indistinguishable from, or assigned to, the unrelated PhaG family, and mannosylerythritol-lipid biosynthesis, a bona fide emulsan locus, and actinobacterial trehalose-mycolate synthesis (the glycolipid class most plausible for the *Actinomycetota* present) were similarly absent. This negative result is not attributable to limited detection sensitivity. Phosphopantetheinyl transferases, without which no NRPS carrier protein can be activated, were abundant in every sample, as were the fatty-acid and sugar precursor pathways that feed lipopeptide and glycolipid synthesis; a community that possesses the activating machinery, the precursor supply, and 69 NRPS clusters, yet essentially no lipopeptide synthetase, is one in which the synthetases are absent rather than merely unresolved by any single method. This conclusion does not rest on the assembly alone: an assembly-independent search of unassembled reads, which cannot be affected by contig fragmentation, recovered lipopeptide-synthetase sequence at a frequency of only about one in 10⁸ reads, and the candidate producing lineage (*Pseudomonas*) was independently confirmed present in the same sample by Kaiju classification. Taken together with the read-level result, the deficit observed at the assembly level is best read as a genuine absence of the synthetases rather than as a product of fragmentation.

Two caveats bound this negative result. First, the screen is only as complete as its reference set: MIBiG curates clustered secondary-metabolite pathways, and several biosurfactant classes plausible for this community, trehalolipids and bacterial glycine lipids among them, are dispersed rather than clustered and are therefore structurally invisible to antiSMASH/MIBiG-based detection; their absence here reflects a limitation of cluster-based methods generally, not evidence specific to this dataset. Second, discriminating closely related enzyme families requires care that gene-name annotation alone cannot provide, as illustrated by the *rhlA*/PhaG case, where sequence sets retrieved under either symbol overlap and correct assignment depends on genomic context and pathway completeness rather than on identity score alone. With these caveats acknowledged, two explanations remain for the absence of secreted biosurfactant biosynthesis: selection against secretion in this habitat, or the taxonomic composition of the community. The lineages that dominate here, salt-in *Halobacteriota* and *Nanohaloarchaeota* in summer and *Flavobacteriaceae*/*Rhodobacteraceae* in winter, are lineages in which lipopeptide and rhamnolipid biosynthesis is rare or unexplored, so their absence is expected on phylogenetic grounds alone and cannot by itself demonstrate active selection. Our data are more consistent with this compositional explanation than with active selection against secretion: biosynthetic capacity scales with genome size in this community (Figure 1B), the streamlined salt-in archaea that dominate summer encode few or no BGCs, and the salt-out bacteria that replace them in winter belong to families in which secreted biosurfactant production is uncommon. Osmotic strategy therefore appears to act as a filter on which biosynthetic capacities are present in the community at all, and not merely on which are expressed, though the present data cannot fully separate this compositional effect from a genuine selective pressure against secretion, and the distinction would require testing biosurfactant capacity in close relatives of these lineages recovered from non-extreme habitats. The near-absence of lipopeptide biosurfactants, combined with strong gene-centred evidence for envelope remodeling and capsule production, supports the view that surface-active adaptation in this hypersaline sediment is primarily cell-retained rather than secreted. This is consistent with the rarity of the lipopeptide core established at the global genomic scale (Ullah et al. 2026) (present in only 0.23% of genomes), and it suggests that earlier reports of widespread biosurfactant production in halophiles, based largely on culture-based activity assays and taxonomic proxies, may have overestimated secreted biosurfactant potential because such methods cannot distinguish cell-associated amphiphiles from genuinely secreted products. Gene-centric, mechanism-specific analysis of the kind applied here offers one way to refine that picture, by separating retained envelope amphiphiles from released biosurfactants at the resolution where the distinction actually matters.

## Conclusion

Genome-resolved metagenomics of seasonally sampled hypersaline sediments shows that this community reorganises between two osmoadaptation strategies, from salt-in haloarchaea in summer to salt-out bacteria relying on compatible-solute biosynthesis in winter, and that its surface-active response under combined cold and osmotic stress is retained at the cell envelope rather than released into the environment. Membrane remodelling, substitution of phospholipids by phosphorus-free ornithine and sulfoquinovosyl lipids, and capsular exopolysaccharide export together account for the amphiphilic signal, while dedicated secreted biosurfactant pathways are absent from the assemblies, from the recovered genomes, from unassembled reads, and from an independent regional genome catalogue.

Two methodological conclusions follow. Taxonomic affiliation substantially overestimates community-level biosurfactant capacity, since the great majority of dominant lineages here carry no such pathway despite belonging to genera reported as producers. Gene-name annotation overestimates it independently, because the committed enzymes of these pathways belong to families whose other members serve primary metabolism, and verification by domain composition reassigned most apparent hits to housekeeping homologues.

These conclusions are bounded. Reference-dependent detection cannot identify a wholly novel synthetase bearing little similarity to characterised references, and the single lipopeptide-initiating cluster recovered here matched no characterised reference and remains functionally unassigned. The design rests on two metagenomes per season, so seasonal contrasts are reported as qualitative patterns cross-validated across independent methods rather than as powered effect sizes. All inferences derive from environmental DNA and describe encoded capacity rather than expressed activity. Metatranscriptomic sampling paired with targeted lipidomics of ornithine and sulfoquinovosyl lipids would establish whether the envelope-remodelling response identified here is actively deployed, and would convert a description of genomic potential into one of realised physiology.

## Author Contributions

***Habib Ullah:*** *Conceptualization, data curation, formal analysis, methodology, software, visualization, and writing original draft. Conducted all bioinformatic analyses*.

***Ramganesh Selvarjan****: Assist in data analysis and manuscript revision*.

***Maggie C. Y. Lau Vetter***: Conceptualization, data interpretation, supervision, and project administration. Provided critical intellectual guidance throughout all stages of the study and reviewed the manuscript.

## Deceleration of Competing Interest

The author declares that there is no conflict of interest.

## Acknowledgments

This work was supported by ANSO (Alliance of International Science Organizations) Scholarship for young talent to H.U., and Discretionary Grant (Institute of Deep-sea Science and Engineering; grant number IDSSE-SJBS-202501), Knowledge Innovation Program of Chinese Academy of Sciences (Institute of Deep-sea Science and Engineering, Chinese Academy of Sciences; grant number Y970011001), Hainan Provincial Funding for Outstanding Talent Team Development (Hainan Provincial Talent Development Bureau; grant number E2500001) to M.C.Y.L.V.

## Data and Code Availability

Raw metagenomic sequencing data generated in this study are deposited in the NCBI Sequence Read Archive under BioProject accession PRJNA1464063 (BioSample accessions SAMN56391696, SAMN56391699, SAMN56391689 and SAMN56391693; SRA accessions SRR38512539 (C3), SRR38512538 (C6), SRR38512537 (H2) and SRR38512536 (H6)); data will be released upon publication. All R scripts and bash scripts used for quality control, assembly, functional annotation, BGC detection, differential abundance analysis, MAG binning, figure generation, and sensitivity analysis are publicly available at GitHub (https://github.com/Habib-Ullah1/QaidamBiosurf-BGC) with a permanently archived version deposited at Zenodo (https://doi.org/10.5281/zenodo.20102918).

